# Epidemiological study on foot-and-mouth disease in small ruminants: sero-prevalence and risk factor assessment in Kenya

**DOI:** 10.1101/2020.05.26.116301

**Authors:** Eunice C. Chepkwony, George C. Gitao, Gerald M. Muchemi, Abraham K. Sangula, Salome W. Kairu-Wanyoike

## Abstract

Foot-and-mouth disease (FMD) is endemic in Kenya affecting cloven-hoofed ruminants. The epidemiology of the disease in small ruminants (SR) is not documented. We carried out a cross-sectional study, the first in Kenya, to estimate the sero-prevalence of FMD in SR and the associated risk factors nationally. Selection of animals to be sampled used a multistage cluster sampling approach. Serum samples totaling 7564 were screened for FMD antibodies of Non-Structural-Proteins using ID Screen^®^ NSP Competition ELISA kit. Data were analysed using Statistical Package for Social Studies Version 20. To identify the risk factors, chi-square and logistic regression analyses were used. The country animal level sero-prevalence was 23.3% (95% CI: 22.3-24.3%) while herd level sero-prevalence was 77.6% (95% CI: 73.9-80.9%). Sero-positivity was significantly higher in the pastoral zone (31.5%) than in the sedentary zone at 14.5% (χ2 =303.2, p<0.05). In the most parsimonious backward fitting logistic multivariable regression, the only risk factors that were significantly positively associated with FMD sero-positivity in SR were multipurpose (OR=1.150; p=0.034) and dairy production types (OR=2.029; p=0.003). Those that were significantly negatively associated with FMD sero-positivity were male sex (OR=0.856; p=0.026), young age (OR=0.601; p=0.037), sedentary production zone (OR=0.471; p<0.001), bringing in of SR (OR=0.838; p=0.004), purchase of SR from market/middlemen (OR=0.877; p=0.049), no interaction with wildlife (OR=0.657; p<0.001), mixed production type (OR=0.701; p=0.016), enclosure of SR day and night (OR=0.515; p=0.001), migratory grazing system (OR=0.807; p=0.047), on-farm watering system (OR=0.724; p=0.002), male-from-another-farm (OR=0.723; p=0.030) and artificial insemination (OR=0.357; p=0.008) breeding methods.

This study showed that there is widespread undetected virus circulation in SR indicated by ubiquitous spatial distribution of significant FMD sero-positivity in the country. The risk factors were mainly husbandry related. Strengthening of risk-based FMD surveillance in carrier SR which pose potential risk of virus transmission to other susceptible species is recommended. Adjustment of husbandry practices to control FMD in SR and in-contact species is suggested. Cross-transmission and more risk factors need to be researched.

## Introduction

Foot and mouth disease (FMD) sometimes referred to as Hoof and mouth disease or Aphthous fever is a highly contagious, transboundary, acute disease caused by foot and mouth disease virus (FMDV) which affects cloven-hoofed domestic ruminants such as cattle, swine, sheep and goats as well as cloven-hoofed wild ruminants [1]. It severely affects livestock production leading to disruption of trade in animals and their products at regional and international level. About 77% of the global livestock population is affected by the disease, mainly in Africa, the Middle East and Asia, and some few areas in South America. This is coupled with the possibility of disease incursion in countries which are currently free. A global strategy for the control of FMD was endorsed in 2012 to minimize the burden of FMD in endemic settings and maintain free status in FMD-free countries [2]. The FMDV is classified into the *Picornaviridae* family and the genus *Apthovirus.* It is a small non-enveloped virus with an icosahedral capsid and a positive sense RNA consisting of a large open reading frame encoding for four structural proteins and ten nonstructural proteins [3]. The FMDV exists in seven immunologically distinct serotypes; A, O, C, Asia 1, SAT 1, SAT 2 and SAT3; all with distinct lineages except SAT 1 and SAT 2 which have unresolved clades [4]. The disease is among the World Organisation for Animal Health (OIE) listed diseases which are transmissible diseases with serious potential to spread across national borders and which require immediate reporting in order to control their spread [2].

The incubation period for foot-and-mouth disease virus is between one and 14 days [5], 3-8 days in small ruminants [6]. The disease is characterized by high fever within two to three days, formation of vesicles and erosions inside the mouth leading to drooling of saliva. Vesicles are also on the nose, teats and when on the feet may rupture and cause lameness. It also causes several months of weight loss in adults and significant deduction in milk production which sometimes fails to return to normal even after recovery. Morbidity rate can be as high as 100% though mortality rate is low in adults but high in neonates due to myocarditis. About 50% of infected ruminants remain asymptomatic carriers in the oropharynx. In cattle the virus can persist for 3-5 years, in sheep up to nine months, in goats up to six months and in the African buffalo up to five years. Pigs do not become carriers [7]. Virus excretion in carrier animals is intermittent and declines over time and the risk of transmission from the African buffalo to cattle exists [8]. Foot and mouth disease in adult sheep and goats is frequently asymptomatic, but can cause high mortality in young animals. Lameness is a significant feature characterized by unwillingness to rise and move [6,9]. The disease can easily be missed unless individual animals are carefully examined for disease lesions. Small ruminants can therefore be responsible for the introduction of FMD into previously disease-free herds [10].The mortality rate in sheep and goats is generally less than 1% in adult animals. Clinical disease in young lambs and kids is characterized by death without the appearance of vesicles, due to heart failure [11].

Although FMD may be suspected based on clinical signs and post-mortem findings, it cannot be differentiated clinically from other vesicular diseases. Confirmation of any suspected FMD case through laboratory tests is therefore essential. The richest source of virus in diagnosis is vesicular fluid or epithelium from fresh lesions. Serum is used for antibody detection where lesions are not fresh and also in epidemiological surveys. Differential diagnosis for FMD in small ruminants includes Peste des petit ruminants (PPR) which can be ruled out by signs of pneumonia and diarrhea, Bluetongue disease (FMD atypical signs are facial oedema and nasal ulceration), Capripox (ruled out by presence of pock lesions), Contagious ecthyma lacks of vesicular stomatitis and lameness which are characteristic in FMD), Pneumonic Pasteurellosis and Contagious Caprine Pleuropneumonia (CCPP) which are characterized by respiratory illness alone [12].

The “gold standard” technique for confirmation of FMD diagnosis is virus isolation [2]. This method is highly sensitive, but lengthy, lasting between one and four days and requires specialised laboratory facilities. The virus neutralization test (VNT) is the “gold standard” technique for detection of antibodies to structural proteins of FMDV and is an approved test for the certification trade of animals and animal products but has variable test sensitivity, is time-consuming, liable to contamination and requires special facilities [13]. Sandwich ELISA is rapid and simple to perform. It is the primary test for FMD diagnosis. The assay depends on serotype-specific polyclonal antibodies prepared in guinea pig and rabbits for the detection of FMDV structural proteins. The test gave 100% specificity and 80% sensitivity in FMDV detection [14]. As demonstrated by [15], detection of the antibodies against the non-structural proteins (NSPs) of FMDV is used for differentiation between infected and vaccinated animals (DIVA) which is of great importance in the control program of FMD. The 3ABC competition antibody ELISA has sensitivity and specificity of the 90% and 99%, respectively, can deliver same-day results when using the short protocol and is routinely applied for general screening for FMD [16,17].

Nucleic acid recognition methods include conventional reverse transcriptase polymerase chain reaction (RT-PCR) assay which is serotyping specific [13]; reverse transcriptase loop-mediated isothermal amplification (RT-LAMP) which is carried out at a constant temperature and does not require a cycler like PCR, is easily performed and can detect FMDV at serotype level in about an hour [18;19]; Multiplex reverse-transcription polymerase chain reaction (mRT-PCR) assay which is a rapid method of high sensitivity of 96.5% for the detection and serotyping of foot and mouth disease virus serotypes [14]. However, the method has not been used extensively, because of factors such as cost, a lack of infrastructure and test complexity. Because of the great sensitivity and specificity of the Real Time RT-PCR assay, it was suggested as the main test for the FMDV detection and is very useful in detecting carrier animals [14].

The FMDV can be transmitted easily through excretions from infected animals (that are newly introduced into a herd; facilities such as buildings or vehicles contaminated with the virus, contaminated materials such as hay, feed, water, milk or biologics; contaminated clothing, footwear, or equipment; raw or inadequately cooked virus-infected meat or other contaminated animal products fed to animals as well as infected aerosols. [2]. Sound biosecurity practices on facilities are required to prevent the introduction and spread of the FMDV into facilities. These practices include livestock and equipment access control, controlled introduction of new animals and materials such as hay into existing herds; regular cleaning and disinfection of livestock pens, buildings, vehicles and equipment; monitoring and reporting of illness as well as appropriate disposal of manure and dead carcasses. Vaccination programmes should target at least 80% of the population in mass vaccinations or designed to target specific animal sub-populations or zones. This should be within the shortest possible time and schedules should be cognizant of maternal immunity [2].

Although the disease causes low mortality in adult animals, the global impact of FMD from direct losses due to reduced production and changes in herd structure as well as indirect losses caused by costs of FMD control, poor access to markets and limited use of improved production technologies is enormous due to the huge numbers of animals affected. Though hard to estimate, the annual economic losses in endemic regions of the world, is between US$ 6.5 and 21 billion and more than US$1.5 billion a year in FMD free countries and zones [20].

Livestock husbandry in developing countries like Kenya is critical for ensuring food security and for poverty alleviation. Goats and sheep (referred to as small ruminants) are normally preferred by farmers compared to large ruminants because of the small space they occupy and less fodder requirement. They important in providing a livelihood, are a source of meat, milk, hides and compost manure as well as an insurance against emergencies [21,22]. In addition, goats have high adaptability to harsh climates which makes them suitable for farmers in marginal areas. The main traditional small ruminant production systems are pastoralism and agro-pastoralism as well as sedentary/mixed systems. Pastoral systems are in the arid and semi-arid climate zones [23] where about 14 million people are dependent on livestock [24]. Agro-pastoralism, is livestock production which is associated with dryland or rain-fed cropping and animals range over short distances. The average sheep and goat herd sizes in pastoralist systems are estimated at 24.9 and 75.2 respectively [25]. Sedentary/mixed systems are found in the semi-arid, sub-humid, humid and highland zones. This farming system is based on livestock but practiced in proximity to, or perhaps in functional association with, other farming systems based on cropping or is a livestock subsystem of integrated crop-livestock farming. The average number of sheep and goats in this system is rarely reported but ranges between three and 10 [23,26,27]. According to the 2009 animal population census, Kenya is home to 17.1 million sheep, 27.7 million goats about 50-57% of which are in the pastoral and agro-pastoral production areas [27, 28]. In Kenya the sheep population is dominated by native breed types such as Red Maasai, Black-head Persian and various types of East African fat tailed sheep. Among goats, the Small East African is most dominant although milk breeds such as the Galla and Toggenberg are also to be found [29].

Commercial dairy and meat goat farming in Kenya is to be found in ranches mainly in the high agricultural potential central and western highlands, central Rift Valley and the coastal region [30]. Besides, some pastoral herds can be large enough to reach commercial proportions [22] Dairy goat farming has become very popular with small scale farmers in urban and densely populated areas of Kenya making a source of living for many people in these areas as goat milk is fast gaining popularity for its nutritional and medicinal value [31]. Meat goats are commonly kept by the pastoral communities and farmers in areas with marginal rainfall and provide a protein delicacy enjoyed by many people. Livestock provides foreign currency to the country and considerably contributes to National Gross Domestic Product (GDP). However, profitability of sheep and goat production is hampered by multifaceted constraints, of which infectious disease is one of the major ones. Foot and Mouth Disease (FMD), Contagious Caprine Pleuropneumonia (CCPP), Pestes de petit ruminants (PPR) and Trypanosomiasis are the major diseases in small ruminants, leading to shortage of milk and meat [32].

Small ruminants (SR) have generally been neglected with regard to their epidemiological role in FMD transmission. This is partly due to the often unapparent nature of the disease in these hosts. Nevertheless, their ability to become carriers represents a reservoir for further infection and spread of disease, and so trade of live sheep and goats present a major risk of entry of FMD to disease-free countries and herds [11]. Table 1 shows the sero-prevalence to FMD in SR and associated risk factors that have been reported in various countries.

**Table 1.** Sero-prevalence to FMD in SR and associated risk factors that have been reported in various countries.

| Country | Sero-prevalence (%) | Associated risk factors | Reference |
| --- | --- | --- | --- |
| Ethiopia | 7.07 in sheep; 7.10 in goats | Agroecology, production system, age | [33] |
| Ethiopia | 4.0-11.0 | Production system, geographic location, species, age, contact with wildlife, season, breed, interaction with other livestock species | [34] |
| India | 23.0 in goats; 12.0 in sheep | Not investigated | [35] |
| Israel | 3.7 | Proximity to herd with outbreak, grazing, herd size | [36] |
| Libya | 13.5 | Not studied | [37] |
| Medina | 27.8 in sheep; 7.9 in goats | Breed | [38] |
| Myanmar | 42.4 | Interaction with infected cattle and pigs, trade area | [39] |
| Nigeria | 41.7 in sheep; 21.8 in goats | Species, geographical location | [40] |
| Pakistan | 21.0 | Age, sex, pregnancy, herd type | [41] |
| Pakistan | 22.8 | Species, sex | [42] |
|  | 25.8 in goats; 14.3 in sheep |  |  |
| Sudan | 14.1 | Not studied | [43] |
| Tanzania | 14.1 in 2015; 39.0 in 2014 | Age, sex, species, geographic location, interaction with other herds | [44] |
| Tanzania (northern) | 48.5 | Age, production system, herd size, acquisition of livestock, wildlife interaction | [45] |
| Uganda | 14 in goats; 22 in sheep | Husbandry practices | [46] |

Therefore, in small ruminants, FMD sero-prevalences in SR of between 3.7% and 48.5% have been reported in various countries. The risk factors that have been associated with FMD sero-positivity in SR in these countries are age, sex, pregnancy, species, breed, herd type, herd size, geographic location, rearing of sheep and goats in contact with infected cattle and pigs, high trade area, proximity to a farm with an FMD outbreak, agro-ecology, season, production system, interaction with other species, interaction with other herds, contact with wildlife and acquisition of livestock.

Foot and mouth disease is endemic in Kenya affecting cattle, sheep, goats, pigs and wild animals such as buffalos and antelopes. Outbreaks are regularly recorded in cattle in Kenya and serotype O has been the most prevalent serotype. Intermittent circulation of FMDV serotypes A, SAT 1 and SAT 2 have also been confirmed in various parts of the country in the last five years [47]. The disease causes widespread outbreaks of clinical disease in cattle although the Government has made efforts to control it. The impact of FMD on national and household economy owing to bans of animals and animal products export to international market outweighs the direct loss due to mortality and morbidity. Studies in pastoral livelihoods in arid and semi-arid areas in Kenya have ranked FMD as second among infectious diseases of livestock and with the highest impact [22,24]. Mortality rate of 40% was reported in lambs in Kenya observed in a serotype O outbreak [48]. There are a large proportion of indigenous and cross-bred small ruminant breeds found in areas where Foot-and-mouth disease frequently occurs.

Despite its huge economic importance, FMD occurrence has not been substantially investigated in small ruminants. Paucity of data cannot provide the real magnitude of the disease in small ruminants in the country. The main control strategies in the country focus on vaccination of cattle. Vaccination against FMD in Kenya is not compulsory. Though SR are also affected by FMD and are herded together with cattle, they are not normally vaccinated. Private farmers are entitled to have their animals vaccinated either by hiring private animal health practitioners or through subsidised government vaccination exercises, if sufficient vaccine is available [49]. Some studies have been carried out on FMD in cattle and buffaloes but no studies on the prevalence and associated risk factors in small ruminants have been done in Kenya.

There are no confirmed reports of distinct FMD outbreaks in indigenous sheep and goats in Kenya. Should SR be involved as reservoirs or amplifiers of virus this would be indicated by a significant incidence of carrier animals and of those showing positive for FMDV antibodies on serological tests. Nevertheless, due to their ability to become carriers and act as reservoirs of infection, this poses major risk of entry of FMD to disease free countries through trade in these animals. There are reports that the silent nature of FMD in small ruminants transmits the virus causing outbreaks by the movement of infected sheep and goats. Good examples include outbreak of FMD in cattle in Tunisia in 1989 which was previously free from FMD and got the infection on importation of sheep and goat from the Middle East [50]. Also in the 2001 FMD outbreak in cattle in the United Kingdom (UK), the role of sheep in the spread of disease was realized. Earlier sheep products such as contaminated frozen lamb from Argentina were blamed for the largest ever recorded type O FMD epidemic in cattle in the UK that occurred in 1967-68 [51]. Thus sheep and goats and their products can play a major role in the transmission of FMD [52].

Sheep and goats form a substantial proportion of the global FMD-susceptible livestock population. However, these species have not been studied with regard to their epidemiological role and significance in the spread of FMD. Unlike cattle, sheep and goats are not included in vaccination control programs in Kenya in spite of their potential infection, although all species are subject to the normal quarantine measures in disease outbreak. Thus, the present study was intended to investigate the sero-prevalence of FMD in small domestic small ruminants in Kenya and to investigate potential risk factors associated with FMD occurrence in these animals. This was achieved through a national cross-sectional study. The results will provide additional knowledge on the role of these ruminants in FMD epidemiology and contribute to development of better Foot and mouth disease control measures in Kenya and globally since FMD is a transboundary disease and also transmitted through export of animals and animal products.

## Materials and Methods

### Description of the Study Area

The study was a cross-sectional study, which targeted the national small ruminant population in Kenya. Kenya is made up of 47 counties (Fig 1). However, the objective was not to primarily measure the sero-prevalence per county but rather per the major small ruminant production zones. The study targeted the second-smallest administrative unit, the sub-location, which was made possible by the fact that the sampling frame of sub-locations was available from the 2009 population and housing census [28].

**Fig 1.**
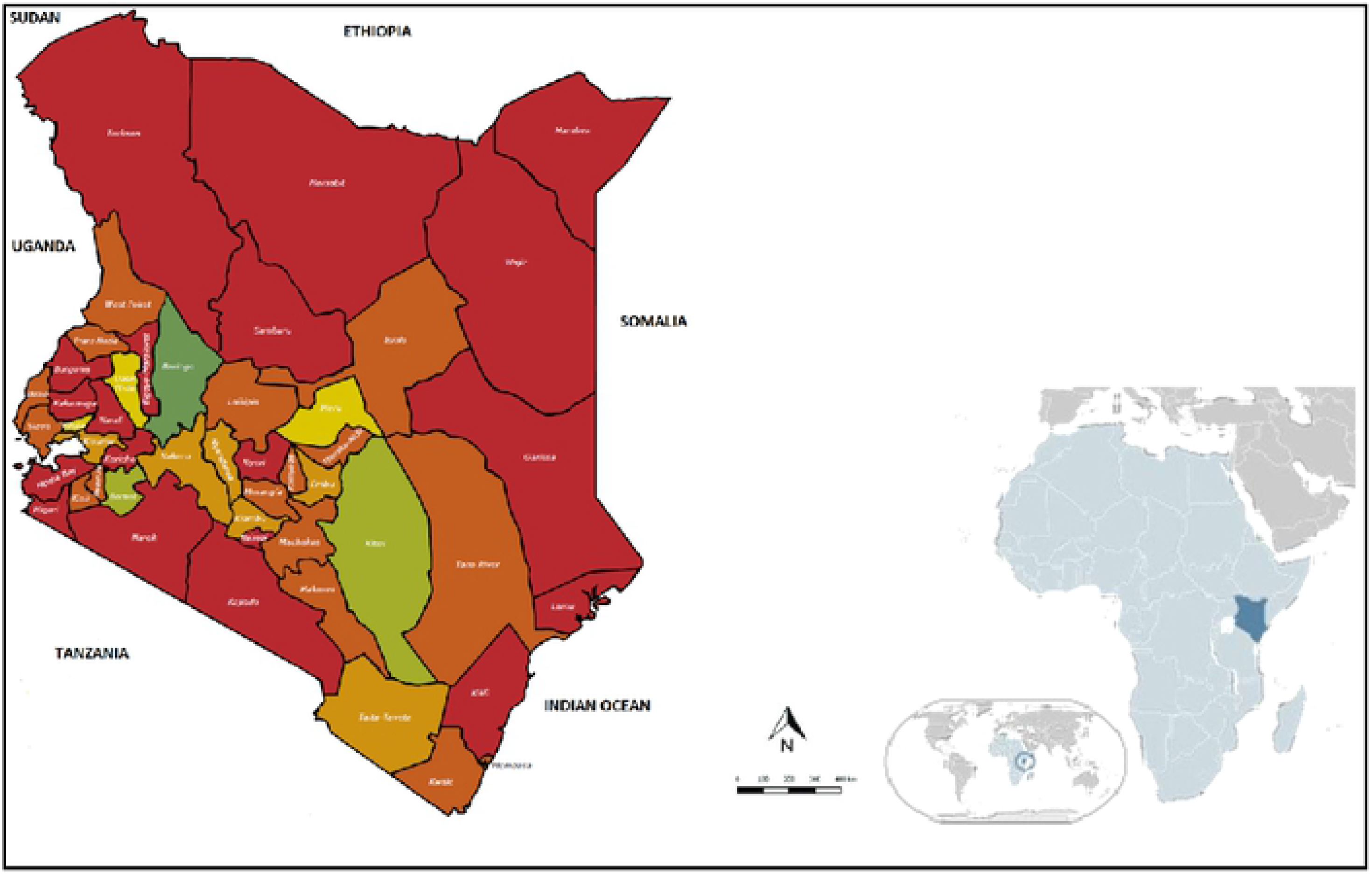
Map of Kenya Counties, 2013 [here].

Kenya lies along the Equator on the extreme Eastern coast of Africa. It lies between 34°E and 42°E and 5°30’N to 4°45’S. The country covers an area of approximately 582,650 square kilometres of which 98% is land while the rest is occupied by water bodies. The human population is 47,564,296 people [53]. Administratively, the country has one unitary national government and 47 county governments. The counties are further divided into sub-counties and wards. Kenya shares common borders with Tanzania to the South (769 Km), Uganda to the West (933 Km), Sudan (232 Km) and Ethiopia (861 Km) to the North and Somalia (682 Km) to the East. It also has a 536 Km coastline on the Indian Ocean.

Broadly, Kenya can be divided into three ecological zones namely humid, semi-arid and arid areas. About 80% of the country is arid and semi-arid (ASAL) while the humid ecosystem occupies the remaining 20% of the country. The humid areas have long rain seasons with heavy down pours reaching 2700mm. The main agricultural activity is mixed crop and livestock farming under sedentary conditions. The semi-arid areas normally experience short rainfall with prolonged drought while arid areas have long cyclic droughts, thus affecting pasture and water availability. Livestock here are kept on a pastoral production system where livestock congregate in search of pasture and water.

### Study design and methods

#### Sample size determination

For the purpose of this study, the country was divided into two zones (strata); the pastoral zone (PZ) and the sedentary zone (SZ) as in Fig 2. Sample size calculation was in two stages and per zone): number of herds to be sampled and then number of animals to be sampled per herd in each herd. The total sample size per stratum was the product of the two. The national sample size was the sum for the two strata. Herd was a group of sheep and goats in a farm/in-contact farms. The number of herds to be sampled was calculated assuming a simple random sample of herds in each stratum independently using Promesa software (http://www.promesa.co.nzl). The assumptions that were made were: number of herds >10,000 in each stratum; confidence level = 95%; accepted margin of error = 5%; expected between herd prevalence = 30%; intra-class correlation coefficient (measure of variation between clusters) = 0.2; design effect = 2[54,55]; test specificity = 99%; test sensitivity = 100% [2]. The calculated sample size was 323 herds per stratum. The minimum sample size for number of animals to be sampled per herd was calculated using Win Episcope 2.0 [56]. This was calculated for a simple random sample, making the following assumptions: expected prevalence of FMD in the herd = 20%; confidence level = 95%; average herd size = 100. This yielded a sample size of 13 animals per herd which was increased to 14 to take care of any possible losses. In each stratum, therefore 323 sub-locations, one village (herd) per sub-location and one household herd per village (or more if necessary to obtain sufficient animals) and 14 animals per herd yielded a sample size of 323×14=4522 samples per stratum and a total of 9044 for the whole country.

**Fig. 2:**
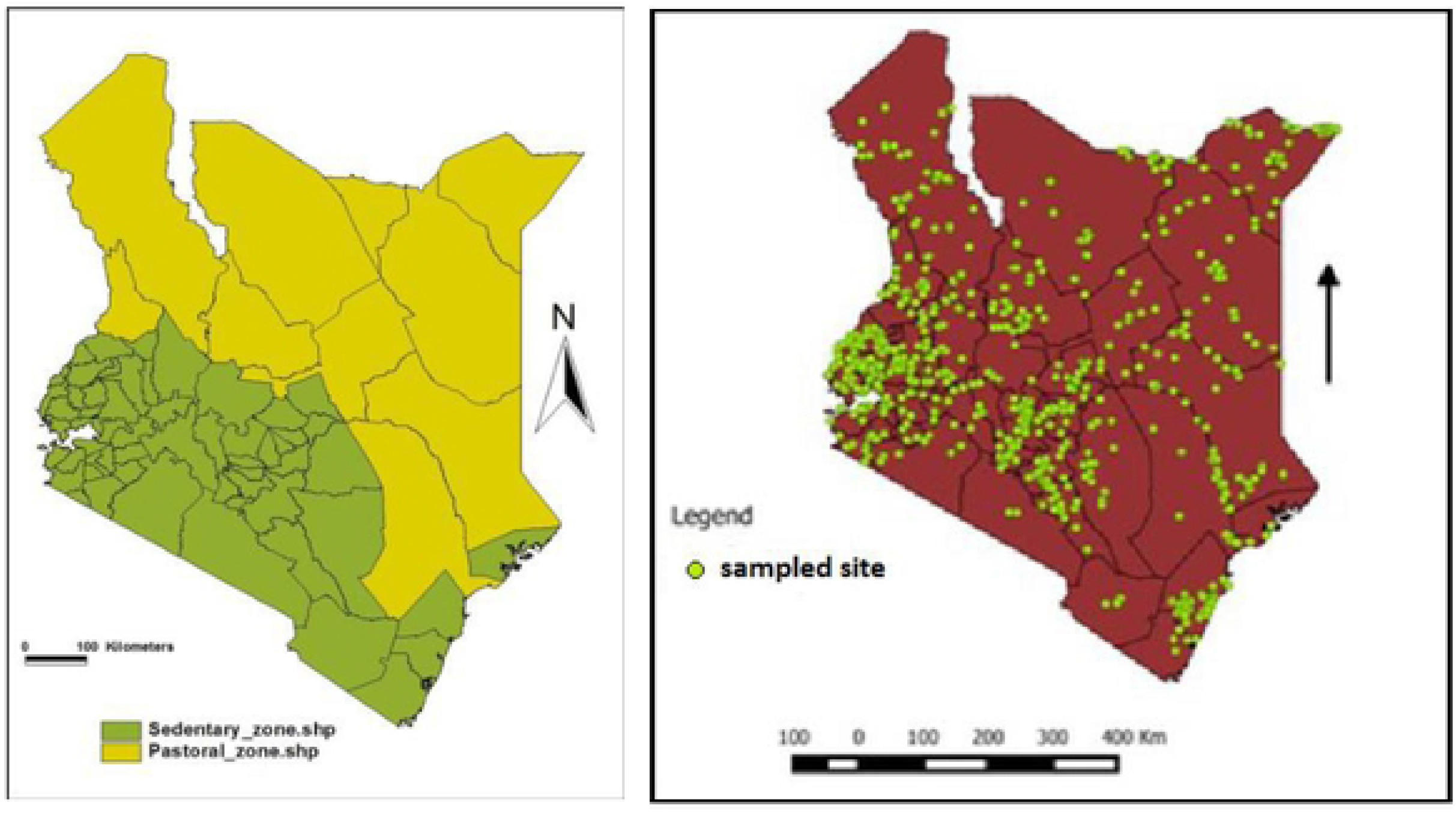
Study zones and selected sampling sites for the cross-sectional survey, Kenya, 2016 [here].

#### Sampling of herds and animals

At the time of survey Kenya had been divided into 47 counties in year 2013 under the new Constitution [57]. However due to available data on the National Census of 2009, the sampling frame were the 6796 Sub-locations and these were used to randomly select the sampling sites. The Country was stratified into two based on livestock production systems – Sedentary and Pastoral with independent sample from each stratum. In each stratum, sample 323 sub-locations. However, sub-locations can be quite large and therefore once in the field the teams obtained the list of villages in the sub-locations and randomly selected villages within the sub-locations (Fig. 2). A herd was then considered as all animals within the village from which the animals sampled were randomly selected. If one herd could yield all the animals required, only that herd was sampled, otherwise additional herds were sampled until the required number to be sampled in the sub-location was reached. Therefore, a multistage cluster sampling was employed to select the animals from the two farming systems. This was by taking sub-location as the first stage, herd (one village per sub-location and one herd from the village) as the second stage and individual animals as the third stage.

#### Data and serum sample collection

Data were collected using questionnaires and from laboratory results while serum samples were collected from eligible herds throughout the country.

##### Serum sample collection

Serum samples were collected by 15 teams of trained laboratory technicians under the supervision of a veterinarian. Sheep and goats aged six months and above were sampled to avoid those with maternal antibodies [58]. Age was determined by examining the dentition of each animal and information from the owner for young animals with no permanent incisors [59]. In the sampling stage animal level variables (biodata) were collected into a sampling form (S3 file) and included species (ovine or caprine), breed, age and the sex of the animal and origin (whether born in herd or brought in). The blood samples were collected from a jugular vein, using 10 ml sterile vacutainer tubes and gauge 21 needles and labeled with a unique identification (county code/sub-location/animal number/sex/age). The samples were then allowed to clot in cool-boxes. Once the blood clot had retracted after 12 to 24 hours the vials were centrifuged in the field laboratories to obtain serum which was placed in two 2ml cryovials (two aliquots) labeled with corresponding identification codes. In areas where laboratories or centrifuges were unavailable serum was separated using sterile disposable pipettes (one per sample) and transferred into the cryovials. Samples were stored at −20°C in field freezers until the end of the sampling period (which was not more than 20 days per team) and were transported on ice using cool-boxes to the Central Veterinary Laboratories (CVL), Kabete, Kenya. At the CVL, the samples were held at −86°C until testing after which they were placed in a serum bank. One aliquot was used to test for the presence or absence of antibodies of Rift Valley Fever (RVF) and Peste de Petits Ruminants (PPR) antibodies according to the objective of the STSD project. The second aliquot was moved to the FMD National Laboratory, Embakasi, Kenya and stored at −20°C until FMD antibodies laboratory investigation.

##### Questionnaire administration

A pre-tested semi-structured questionnaire (S1 file) was administered in-person by trained enumerators to owners of sampled herds following the guidelines (S2 file) at the time of sample collection for collection of herd level variables. Herd-level variables were production zone, whether the herd owners brought in animals in the last one year, whether the herd owners purchased animals from the market/middlemen, interaction with wildlife, production type, production system, housing type, grazing system, watering system, breeding method and altitude. Also based on Geographic Positioning System (GPS) technology, GPS coordinates and elevations were recorded for each herd location and this information was recorded in each questionnaire form which was labeled with the unique herd identification number.

#### Laboratory sample analysis

The serum samples were analyzed by ID Screen FMD NSP competition foot and mouth disease virus 3 ABC- ELISA (ID Screen®FMD NSP Competition) kit to detect specific antibodies against the non-structural protein of Foot and Mouth Disease virus (FMDV NSP). On the seropositivity/negativity to FMDV antibodies the outcome variables were categorised based on the on the results of the 3ABC blocking enzyme-linked immunosorbent assay. Briefly in this procedure, samples were exposed to non-structural FMDV antigen (NSP 3 ABC) coated wells on micro titer plates; samples to be tested and the controls were added to the microwells; anti-NSP-antibodies if present form an antigen-antibody complex which masks the virus epitopes; anti-horseradish peroxidase (HRP) conjugate is added subsequently to the microwells and it fixes the remaining free epitopes forming an antigen-conjugate HRP-complex. The excess conjugate is removed by washing and the substrate solution; 3,3,5,5 - tetramethylbenzidine (TMB) is added. The resulting coloration depends on the quantity of specific antibodies present in the sample being tested. In the absence of antibodies a blue solution appears which becomes yellow after addition of stop solution. In the presence of antibodies no coloration appears. Within 15 min in the dark, the result was read by micro plate spectro-photometer at 450 nm optical wavelength. The diagnostic relevance of the result was obtained by comparing the OD which develops in wells containing the samples with the OD from the wells containing the positive control as it was read by the ELISA reader. To validate the test the mean OD value of negative control is greater than 0.7 while mean OD for positive control is less than 0.3. Sera with competition percentage ratio for the test sample and negative control (S/N %) less than or equal to 50% are considered positive and S/N% greater than 50% are considered negative. On herd level sero-prevalence a herd was considered as positive if one or more animals in the herd were seropositive.

#### Data management and analysis

Individual animal laboratory data generated during testing along with individual animal biodata data obtained during sample collection (species, breed, sex, age, origin) were entered in Microsoft Excel 2010 spreadsheet (S4 and S5 files). Questionnaire data which included mainly herd data (production zone, whether animals were brought into the herd, whether animals were purchased from markets and middlemen, wildlife interaction, production type, production system, housing system, grazing system, watering system, breeding method and location altitude) were entered in Microsoft Access 2010. Both data sets were then brought together in a Microsoft Excel Spreadsheet, cleaned and coded before being exported to Statistical Package for Social Science (SPSS) Version 20 for analysis. Analysis included descriptive analysis of the variables to generate means, medians, proportions and confidence intervals. Descriptive statistics were also generated for the sero-prevalence in the two different production zones (pastoral and sedentary), for each county and for the other risk factors. Apparent prevalence was calculated using equation 1 while true prevalence was calculated using equation 2

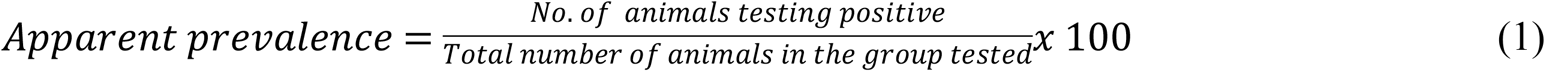

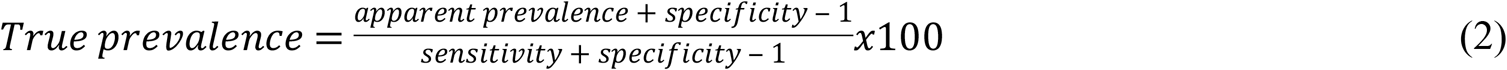

Chi-squared test as recommended by Campbell [60] and Richardson [61] was used to compare proportions while the confidence intervals of the proportions were calculated using the method recommended by Altman et al. [62]. The test of crude association between risk factors (both individual animal and herd level) and FMD sero-positivity was done using chi-square test. Bivariable and multivariable logistic regression analyses were used to test the strength of association between the risk factors and FMD sero-positivity, making use of Odds Ratio (OR) and p values. The cut-off p value was 0.05.

Coding took into consideration the regression analyses such that the lowest code (0) was for the factor that was considered as the reference. The reference was the factor which exhibited the highest proportion/Wald statistic https://www.theanalysisfactor.com/strategies-dummy-coding/. Risk factors from the bivariable regression model entering multivariable regression analysis were not necessarily those with p≤ 0.05, but p≤0.2, so that any variables that became significant once confounding was removed through multivariable regression analysis were not missed out [63]. Test for collinearity of the variables was by testing for correlation. Simple correlation coefficients for pairs of independent variables are determined and a value of 0.8 or more implies severe collinearity among the affected independent variables and one has to be dropped [64]. In the interpretation of results in all the analysis, confidence level was kept at 95% and ρ≤0.05 was set for significance. If the probability value (p value) was less than or equal to 0.05, then the result was considered as statistically significant. Ultimate strength of association between sero-positivity and risk factors was through interpretation of the OR obtained in multivariable regression analysis. The interpretation of odds ratios less than one were after obtaining their inverse [65].

The goodness of fit tests used for the regression models were the Omnibus test of model coefficients and Hosmer and Lemeshow test. The omnibus test looks for improvement of the new model (with explanatory variables included) over the baseline model. It uses chi-square tests to scan for significant difference between the Log-likelihoods (specifically the (−2Log Likelihood, −2LL) of the baseline model and the new model. If there is a significant reduction in −2LL in the new model compared to the baseline then it suggests that the new model is an improvement [66]. The Hosmer and Lemeshow statistic is calculated using −2LL and produces a p-value based on a chi-square distribution. It tests the null hypothesis that the model is a good enough fit for the data. The null hypothesis is rejected if p<0.05 [67].

The results of our study have been presented mainly in tables and figures and interpreted in text. Although the bivariable regression was carried out for all risk factors (individual animal and herd level) together, the results are in two tables to avoid too large a table.

#### Ethics

The research approval for the study was obtained from the Kenya National Commission For Science, Technology and Innovation (NACOSTI). Each owner of a herd selected for sampling provided verbal consent, once the objectives of the study were explained. Herds whose owners did not consent were replaced with the next herd in the random sample list. Other approvals required for the study were obtained from the State Department of Livestock at national level and from the respective County governments.

## Results

The cross-sectional study was carried out from August to September 2016 cross-nationally. In the study, 898 herds (herd was used to mean a group of sheep and goats) were sampled yielding 8201 samples. However, only 7564 samples from 872 herds were available for testing for FMD sero-prevalence. Sheep samples were 2560 (33.8%) while goat samples were 5004 (66.2%). Of these 3909 (51.7%) were from the PZ and 3655 (48.3%) were from the SZ. Of the 44 Counties investigated, 11 (25.0%) were from PZ and 33 (75.0%) were from the SZ. Though slightly lower than the calculated sample size, due to sample loss and lack of usability of some samples for the test (16%), these samples were deemed sufficient for determination of the sero-prevalence of FMD in the SR herds given that there was high design effect (2) consideration and provision for sample loss in sample size calculation.

### Animal and herd level descriptive statistics

Table 2 shows the number of sheep and goats in the sampled herds in the PZ and SZ. Therefore, the mean herd size in the PZ was about ten times that in the SZ.

**Table 2.** Number of sheep and goats in the herds sampled in the pastoral and sedentary zones, Kenya, 2016.

| **Pastoral Zone (PZ)** | **Minimum** | **Maximum** | **Sum** | **Mean** | **Std. Deviation** | **95% CI of mean** |
| --- | --- | --- | --- | --- | --- | --- |
| Male<1yr | 1 | 330 | 9,761 | 12.93 | 25.33 | 9.66-14.44 |
| Female<1yr | 1 | 500 | 15,600 | 18.39 | 40.01 | 14.87-22.82 |
| Male≥1yr | 1 | 250 | 12,539 | 14.09 | 23.19 | 12.02-16.60 |
| Female≥1yr | 1 | 750 | 42,696 | 64.68 | 98.94 | 55.53-74.86 |
| Overall Mean | 1 | 458 | 20,149 | 27.50 |  |  |
| **Sedentary zone (SZ)** |  |  |  |  |  |  |
| Male<1yr | 1 | 100 | 1,523 | 1.67 | 5.15 | 1.30-2.20 |
| Female<1yr | 1 | 25 | 1,545 | 1.72 | 2.90 | 1.48-1.98 |
| Male≥1yr | 1 | 300 | 2,409 | 2.82 | 14.51 | 1.90-4.26 |
| Female≥1yr | 1 | 160 | 5,561 | 4.61 | 11.90 | 3.65-5.74 |
| Overall Mean | 1 | 146 | 2,760 | 2.70 |  |  |

S1 Table presents the individual animal variable descriptive statistics in both the PZ and SZ and overall. Thus about two-thirds (63.2-68.9%) of the SR sampled were of caprine species in both zones and in total. Nearly half of the animals (43.8%) were of local breed in both zones and in total. However a large proportion of animals (30.5-40.6%) in both zones and overall had their breeds unidentified. Over three-quarters (77.0-79.9%) of the SR sampled in both zones and in total were female.

S2 Table presents the herd level variable descriptive statistics in both the PZ and SZ and overall. Although these were herd level variables, the numbers are of actual number of animals involved as most analyses in the study were at individual animal level. Regarding general practices within the herds, only about a third of animals were in herds (35.5-37.8%) where SR were brought into the herds in the last one year in both the PZ and SZ and overall. While about half (55.7%) of the animals in the PZ were in herds which purchased SR from markets or middlemen, only a small proportion (17.4%) of animals in the SZ were from herds which purchased so. Ultimately, nearly one third of animals (37.2% were from herds that purchased so in the overall. A large proportion of animals (83.0%) in the PZ were from herds which had interaction with wildlife. In the sedentary zone, about one third (28.8%) were from herds with such interaction while for another about one third (37.2%), it was not clear whether the herds they came from had such interaction or not.

With regard to SR husbandry practices, in both zones and overall, majority (83.2-88.8%) of animals were in herds where the production type was meat or multipurpose. In the PZ, majority of animals (83.5%) were in herds which were under pastoral and agro-pastoral production systems hence the categorization as PZ. On the other hand, majority of animals (78.0%) in the SZ were in herds under sedentary/mixed production systems hence the categorization SZ. Overall most animals (89.4%) were from herds with pastoral, agro-pastoral and sedentary/mixed production systems. About three quarters of animals (75.5-78.3%) in both zones and overall were from herds that were enclosed at night in both zones and overall. However, a significant proportion of animals (16.2-20.3%) in all three scenarios were from herds that were not enclosed at all. Majority of animals in the PZ were in herds which had communal (41.2%) or mixed (27.4%) grazing systems. Only 19.0% were under migratory (nomadic) grazing system. Similarly in the SZ, a substantial proportion of animals (30.5%) were from herds with communal grazing system but more (49.2%) were from herds with fenced grazing systems. Overall, 80.8% of animals were in herds with communal, fenced and mixed grazing systems. Majority of animals (90.3%) in the PZ were from herds where shared watering was practiced. In the SZ, most animals (60.7%) were in herds with on-farm watering system although a substantial proportion (27.7%) was also in herds where shared watering was practiced. Overall, the scenario on watering was mainly shared (60.0%) but also on-farm (32.1%) watering system. The breeding method in the PZ in the herds where most animals (52.7%) were contained was own-male but also mixed and common-use methods (40.1%). In the SZ, majority of animals were in herds where the breeding method used was own-male (64.8%) but also male-from-another-farm (10.5%). Overall, the scenario on breeding method was mainly own-male (58.6%) but also mixed and common-use-male (27.0%). While animals in the PZ were mainly (80.1%) in herds at low altitude (≤1500m above sea level), animals in the SZ were nearly equally distributed in herds at low altitude and at high altitude (>1500m above sea level) at 48% and 50% respectively.

### Sero-prevalence of FMD in small ruminants

Sero-prevalence of a total of 7564 sera from the whole country was determined for the presence of non-structural FMDV protein (antibodies). The overall sero-prevalence of FMD in small ruminants was 23.3% (95% CI: 22.3 - 24.3%). The prevalence rate was significantly higher (χ2 = 305.55, ρ<0.001) in the PZ at 31.5% (95% CI: 29.8-32.7%) compared to that in the SZ which had a prevalence of 14.5% (95% CI: 13.4-15.7%). The sero-prevalence per County is in the S3 Table. Variations in spatial distributions of FMD sero-prevalence were observed across the country with the highest apparent sero-prevalence recorded in Mandera at 64.5%, Kilifi 49.1%, Lamu 42.9%, Kajiado 38.6%, West Pokot 36.0%, Garissa 34.5%, Turkana 28.9% and Tana River 26.0%. Notably these counties had higher seroprevalence than the national average. Many counties in SZ had low seroprevalence rate of less than 10.0% namely Embu, Kisii, Nakuru, Elgeiyo-Marakwet, Kiambu, Bungoma, Kirinyaga, Vihiga and Muranga counties. Many other counties in sedentary zone had seroprevalence rate above 10.0% but lower than the national level of 23.3%. Mombasa and Nyamira showed sero-negativity but the number of samples tested was small (5 and 14 respectively) to give any meaningful interpretation. The test used (NSP ELISA) has 100% sensitivity and 99% specificity [2] and the true prevalence was therefore lower than the apparent sero-prevalence.

The distribution of sites where samples tested positive is as in Fig 3. Thus the distribution of FMD sero-positives among SR was near ubiquitous with nearly every county registering some positives. The positive sites appear to be more concentrated in the SZ but this was because more sites were needed in the SZ to yield the required sample size since herds there are smaller.

**Fig 2.**
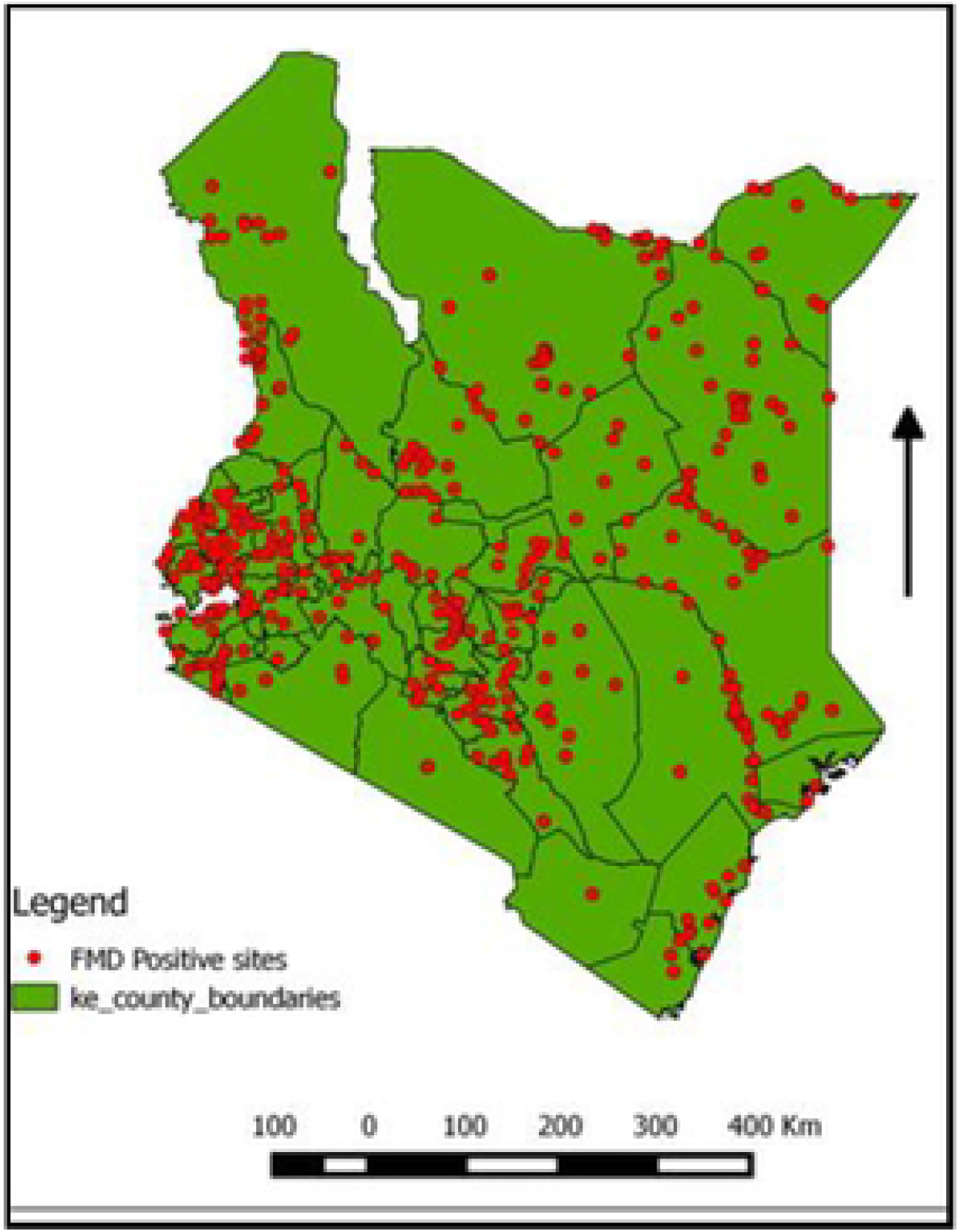
Map of sites where samples tested positive for FMD, Kenya, 2017 [Here].

Pearson’s correlation of all the risk factors showed a strong correlation between county type and production zone (r=0.992; p<0.001). For this reason, only production zone was retained in the group of risk factors (among others) for FMD sero-positivity risk factor analysis. The sero-positivity per individual animal risk factor was as in Table 3.

**Table 3.** Sero-positivity per individual animal risk factor, Kenya, 2016.

| Variable | Variable Code | Total tested | Positive | Negative | % Positive | 95%CI of % positive |
| --- | --- | --- | --- | --- | --- | --- |
| **Species** |  |  |  |  |  |  |
| Caprine | 0 | 5004 | 1202 | 3802 | 24.0 | 22.9-25.2 |
| Ovine | 1 | 2560 | 560 | 2000 | 21.9 | 20.3-23.5 |
| **Breed** |  |  |  |  |  |  |
| Local | 0 | 3310 | 807 | 2503 | 24.4 | 22.9-25.9 |
| Cross-breed | 1 | 861 | 207 | 654 | 24.0 | 21.2-27.1 |
| Exotic | 2 | 694 | 123 | 571 | 17.7 | 15.0-20.8 |
| Unidentified<sup>o</sup> | 3 | 2699 | 25 | 2074 | 0.9 | 0.6-1.4 |
| **Sex** |  |  |  |  |  |  |
| Female | 0 | 5922 | 1403 | 4519 | 23.7 | 22.6-24.8 |
| Male | 1 | 1636 | 356 | 1280 | 21.8 | 19.8-23.9 |
| Unidentified | 2 | 6 | 3 | 3 | 50.0 | 14.0-86.1 |
| **Age** |  |  |  |  |  |  |
| Mature (>1 year) | 0 | 7335 | 1731 | 5604 | 23.6 | 22.6-24.6 |
| Young ($\leq 1$ year) | 1 | 198 | 20 | 178 | 10.1 | 6.4-15.4 |
| Unidentified | 2 | 31 | 20 | 11 | 35.5 | 19.8-54.6 |
| **Origin** |  |  |  |  |  |  |
| Born in herd | 0 | 6273 | 1474 | 4799 | 23.5 | 22.5-24.6 |
| Brought in | 1 | 317 | 54 | 263 | 17.0 | 13.2-21.7 |
| Unidentified | 2 | 974 | 234 | 740 | 24.0 | 21.4-26.9 |

<sup>o</sup>Unidentified means that the variable was not indicated for those samples

Thus at individual animal level, the sero-positivity of FMD in caprine (goats) compared to that in ovine (sheep) was higher but this was not statistically significant. That for exotic breeds was significantly lower than that for local breeds (χ2=14.43; p<0.001) and cross breeds (χ2=9.13; p=0.003). Similarly, the sero-prevalence in mature animals was higher than in young animals and this was statistically significant (χ2=19.69; p<0.001), while that in animals that were born in the herd was significantly higher than that of animals that were brought in (χ2=7.16; p=0.008). However, that in female animals was higher than in male animals but this was not statistically significant (χ2= 2.59 and p=0.108).

The between herd prevalence, a measure of sero-prevalence in herds where at least one animal in a herd tested positive, for all the 872 herds tested was 77.6% (95% CI: 73.9-80.9), which was significantly higher than animal level sero-prevalence (23.3% (95% CI: 22.3 - 24.3). The sero-positivity per herd risk factor was as in Table 4. With regard to herd level sero-positivity for FMD in SR, in the PZ was significantly higher than in the SZ (χ^2^ = 295.15; p<0.001).

**Table 4:** Sero-positivity for FMD per herd risk factor, Kenya, 2016.

| Variable | Variable Code | Total tested | Positive | Negative | % Positive | 95%CI of % positive |
| --- | --- | --- | --- | --- | --- | --- |
| **Production Zone** |  |  |  |  |  |  |
| Pastoral | 0 | 3909 | 1220 | 2689 | 31.2 | 29.8-32.7 |
| Sedentary | 1 | 3655 | 531 | 3124 | 14.7 | 13.4-15.7 |
| **Herds brought in SR?** |  |  |  |  |  |  |
| No | 0 | 4767 | 1160 | 3607 | 24.3 | 23.1-25.6 |
| Yes | 1 | 2769 | 598 | 2171 | 21.6 | 20.1-23.2 |
| Unidentified<sup>o</sup> |  | 28 | 4 | 24 | 14.3 | 4.7-33.6 |
| **Buy SR from market or middlemen?** |  |  |  |  |  |  |
| No | 0 | 4753 | 1023 | 3730 | 21.5 | 20.4-22.7 |
| Yes | 1 | 2811 | 739 | 2072 | 26.3 | 24.7-28.0 |
| **SR interaction with wildlife** |  |  |  |  |  |  |
| Yes | 0 | 4299 | 1201 | 3098 | 27.9 | 26.6-29.3 |
| No | 1 | 1451 | 183 | 1268 | 12.6 | 11.0-14.5 |
| Unidentified | 2 | 1814 | 378 | 1436 | 20.8 | 19.0-22.8 |
| **SR Production type** |  |  |  |  |  |  |
| Meat | 0 | 3341 | 700 | 2641 | 21.0 | 19.6-22.4 |
| Multipurpose | 1 | 3170 | 816 | 2354 | 25.7 | 24.2-27.3 |
| Mixed | 2 | 366 | 70 | 296 | 19.1 | 15.3-23.6 |
| Dairy | 3 | 190 | 31 | 159 | 16.3 | 11.5-22.5 |
| Unidentified | 4 | 497 | 145 | 352 | 29.2 | 25.3-33.4 |
| **SR Production System** |  |  |  |  |  |  |
| Sedentary/mixed | 0 | 3115 | 463 | 2652 | 14.9 | 13.6-16.2 |
| Pastoral | 1 | 2593 | 799 | 1794 | 30.8 | 29.0-32.6 |
| Agro-pastoral | 2 | 1048 | 305 | 743 | 29.1 | 26.4-32.0 |
| Multiple | 3 | 166 | 37 | 129 | 22.3 | 16.4-29.5 |
| Unidentified | 4 | 642 | 158 | 484 | 24.6 | 21.4-28.2 |
| **SR Housing** |  |  |  |  |  |  |
| Enclosed at night | 0 | 5821 | 1377 | 4444 | 23.7 | 22.6-24.8 |
| None | 1 | 1386 | 346 | 1040 | 25.0 | 22.7-27.3 |
| Enclosed day and night | 2 | 357 | 39 | 318 | 10.9 | 8.0-14.7 |
| **SR grazing** |  |  |  |  |  |  |
| Communal | 0 | 2724 | 713 | 2011 | 26.2 | 24.5-27.9 |
| Fenced | 1 | 2117 | 335 | 1782 | 15.8 | 14.3-17.5 |
| Mixed | 2 | 1268 | 351 | 917 | 27.7 | 25.3-30.3 |
| Migratory | 3 | 786 | 232 | 554 | 29.5 | 26.4-32.9 |
| Unidentified | 4 | 358 | 96 | 262 | 26.8 | 22.4-31.8 |
| Zero-grazing | 5 | 311 | 35 | 276 | 11.3 | 8.1-15.4 |
| **SR watering** |  |  |  |  |  |  |
| Shared | 0 | 4540 | 1290 | 3250 | 28.4 | 27.1-29.8 |
| On-farm | 1 | 2425 | 330 | 2095 | 13.6 | 12.3-15.1 |
| Unidentified | 2 | 455 | 124 | 331 | 27.3 | 23.3-31.6 |
| Mixed | 3 | 144 | 18 | 126 | 12.5 | 7.8-19.3 |
| **SR breeding method** |  |  |  |  |  |  |
| Own male | 0 | 4429 | 1020 | 3409 | 23.0 | 21.8-24.3 |
| Mixed | 1 | 1244 | 325 | 919 | 26.1 | 23.7-28.7 |
| Common-use male | 2 | 800 | 215 | 585 | 26.9 | 23.9-30.1 |
| Unidentified | 3 | 578 | 131 | 447 | 22.7 | 19.4-26.3 |
| Male from another farm | 4 | 446 | 63 | 383 | 14.1 | 11.1-17.8 |
| Artificial insemination | 5 | 67 | 8 | 59 | 11.9 | 5.7-22.7 |
| **SR location elevation** |  |  |  |  |  |  |
| ≤1500m | 0 | 4884 | 1277 | 3607 | 26.2 | 24.9-27.4 |
| >1500m | 1 | 2469 | 419 | 2050 | 17.0 | 15.5-18.5 |
| Unidentified | 2 | 211 | 66 | 145 | 31.3 | 25.2-38.1 |

<sup>¶</sup>Unidentified means that the variable was not indicated for those samples

Unexpectedly, herds that did not bring in small ruminants had a higher sero-positivity than those that brought in small ruminants although this was not statistically significant (χ^2^=7.14; p=0.008). The sero-positivity of herds in which animals were bought from the market or middlemen was significantly higher than in those herds where this was not the case (χ^2^=22.78; p<0.001). Similarly, herds which had wildlife interaction had significantly higher sero-positivity than those without such interaction (χ^2^ =139.05; p<0.001). The sero-positivity of herds at low altitude (≤1500m above sea level was significantly higher (χ^2^ =78.10; p=0.001) than that of herds at higher altitude (>1500m above sea level).

Regarding animal husbandry, multipurpose production type herds had significantly higher sero-positivity than herds with all other surveyed production types while there was no significant difference in sero-positivity among all other production types. The pastoral production system showed significantly higher sero-positivity than sedentary/mixed production system (χ^2^ =207.61; p<0.001). The difference between the pastoral production system and other production systems (agro-pastoral, multiple) was not statistically significant (p>0.05). Herds in which animals were not enclosed or enclosed only at night had a significantly higher sero-positivity than herds which were enclosed by day and by night (χ^2^ =32.75; p<0.001 and χ^2^=31.15; p<0.001 respectively). Communal, mixed and migratory grazed herds did not show significant difference in sero-positivity. However, all showed a significant difference in sero-positivity with fenced herds and zero-grazed herds. There was no significant difference in sero-positivity between fenced and zero-grazed herds (χ^2^ =4.25; p=0.039). Small ruminant herds that had shared watering had significantly higher sero-positivity than those with on farm watering and mixed type watering (χ^2^ =194.02; p<0.001; χ^2^ =17.76; p<0.001respectively). There was no significant difference between on farm and mixed watering methods. Regarding breeding methods, unexpected results were observed. There was no significant difference in sero-positivity with regard to use of own-male, common-use-male and mixed breeding methods. There was however, significant difference in sero-positivity between the aforementioned breeding methods and male-from-another-farm. Expectedly there was a significant difference in sero-prevalence is herds with aforementioned breeding methods and artificial insemination which had the lowest sero-prevalence. The difference in sero-positivity between herds with male-from-another-farm and artificial insemination was statistically insignificant.

### Association between FMD sero-positivity and selected risk factors

The crude association between sero-positivity and the selected risk factors was determined using chi-square. The Chi-square results are in Table 5. The only risk factor at individual animal level and herd level that did not show statistically significant association with sero-positivity to FMD was sex of the small ruminant.

**Table 5.** Crude association between sero-positivity and risk factors, Kenya, 2016.

| Variable | Chi-square | p |
| --- | --- | --- |
| Species | 4.245 | 0.039 |
| Breed | 14.538 | 0.002 |
| Sex | 5.072 | 0.079 |
| Age | 22.248 | <0.001 |
| Origin | 7.387 | 0.025 |
| Production zone | 294.252 | <0.001 |
| Herds brought in SR? | 8.624 | 0.013 |
| Buy SR from market or middlemen? | 22.458 | <0.001 |
| SR<sup>†</sup> interaction with wildlife | 150.644 | <0.001 |
| SR Production type | 39.240 | <0.001 |
| SR Production System | 226.470 | <0.001 |
| SR Housing | 33.160 | <0.001 |
| SR grazing | 137.167 | <0.001 |
| SR watering | 207.309 | <0.001 |
| SR breeding method | 37.439 | <0.001 |
| SR location elevation | 85.027 | <0.001 |

<sup>†</sup>SR means small ruminant

According to the bivariable regression model (S4 Table), there were no individual animal risk factors that showed positive association with sero-positivity for FMD in small ruminants. All the risk factors: species (ovine), breed (cross, exotic), sex (male), age (young) and origin (brought in) showed negative association. Ultimately, the statistically significant association was with species (OR=0.886; 95%CI: 0.790-0.993; p=0.037), exotic breed (OR=0.668; 95%CI: 0.541-0.825; p<0.001), age (OR=0.364; 95%CI: 0.228-0.579; p<0.001) and origin (OR=0.668; 95%CI: 0.496-0.901; p=0.008).

The bivariable regression model for herd level risk factors for FMD in small ruminants is in S5 Table. According to this model, the risk factors which showed statistically significant positive association with sero-prevalence for FMD in SR compared to their respective references were purchase from market/middlemen (OR=1.300; 95%CI: 1.166-1.450; p<0.001); multipurpose production type (OR=1.308; 95%CI: 1.165-1.468; p<0.001), pastoral production system (OR=2.551; 95%CI: 2.242-2.903; p<0.001), agro-pastoral production system (OR=2.351; 95%CI: 1.992-2.775; p<0.001) and multiple production system (OR=1.643; 95%CI: 1.125-2.399; p<0.010), mixed breeding method (OR=1.182; 95%CI: 1.023-1.366; p=0.023) and common-use-male breeding method (OR=1.228; 95%CI: 1.035-1.458; p=0.019). Conversely, the risk factors which showed statistically significant negative association with sero-prevalence for FMD in SR were sedentary production zone (OR=0.377; 95%CI: 0.336-0.422; p<0.001), herd brought in SR (OR=0.875; 95%CI: 0.766-0.958; p=0.007), no interaction with wildlife (OR=0.372; 95%CI: 0.314-0.441; p<0.001), enclosure of animals day and night (OR=0.396; 95%CI: 0.282-0.555; p<0.001), fenced grazing type (OR=0.530; 95%CI: 0.457-0.613-1.468; p<0.001) and zero-grazing (OR=0.358; 95%CI: 0.249-0.514; p<0.001).

All the 16 variables considered in the bivariable regression model were retained in the multivariable regression analysis because at least one category in each had p≤0.2. After controlling for confounding, the most parsimonious backward fitting logistic multivariable regression model was as in Table 6. In the final model obtained, the omnibus test result was χ2=465.04 df=35 p<0.001which demonstrated a significant change in the coefficients of most variables compared to the base model with the conclusion that the model was a good fit for the data. With the Hosmer and Lemeshow test p=0.712 and therefore the null hypothesis was rejected leading to a result that the model was a good fit for the data. These goodness-of fit test results were considered reliable given the neither too small nor too big sample size of 7564. In this model, the only risk factors that were significantly positively associated with sero-positivity of FMD in SR were multipurpose and dairy production types. Those that were significantly negatively associated were sex (male), young age, sedentary production zone, bringing in of SR, purchase of SR from market/middlemen, no interaction with wildlife, mixed SR production type, enclosure of SR day and night and migratory grazing system, on-farm watering system, male-from-another farm and artificial insemination breeding methods.

**Table .:** Multivariable regression model of risk factors for FMD sero-positivity in small ruminants, Kenya, 2016.

| Risk factor | Variable | B | S.E. | Wald | p | OR | 95%CI of OR |
| --- | --- | --- | --- | --- | --- | --- | --- |
| Sex | Female | Ref |  |  |  |  |  |
|  | Male | -0.155 | 0.070 | 4.945 | 0.026 | 0.856 | 0.747-0.982 |
|  | Unidentified | 0.710 | 0.866 | 0.673 | 0.412 | 2.034 | 0.373-11.096 |
| Age | Mature | Ref |  |  |  |  |  |
|  | Young | -0.509 | 0.244 | 4.352 | 0.037 | 0.601 | 0.373-0.970 |
|  | Unidentified | 0.448 | 0.409 | 1.197 | 0.274 | 1.565 | 0.702-3.492 |
| Origin | Born in herd | Ref |  |  |  |  |  |
|  | Brought in | -0.016 | 0.161 | 0.010 | 0.921 | 0.984 | 0.717-1.350 |
|  | Unidentified | -0.244 | 0.095 | 6.529 | 0.011 | 0.784 | 0.650-0.945 |
| Production Zone | Pastoral | Ref |  |  |  |  |  |
|  | Sedentary | -.754 | .088 | 73.704 | <0.001 | 0.471 | 0.396-0.559 |
| Brought in SR | No | Ref |  |  |  |  |  |
|  | Yes | -0.176 | 0.062 | 8.089 | 0.004 | 0.838 | 0.742-0.947 |
|  | Unidentified | -0.718 | 0.554 | 1.680 | 0.195 | 0.488 | 0.165-1.445 |
| Buy SR from market or middlemen? | No | Ref |  |  |  |  |  |
|  | Yes | -0.131 | 0.067 | 3.887 | 0.049 | 0.877 | 0.769-0.999 |
| SR interaction with wildlife? | Yes |  |  |  |  |  |  |
|  | No | -0.420 | 0.105 | 15.919 | <0.001 | 0.657 | 0.535-0.808 |
|  | Unidentified | 0.122 | 0.086 | 2.007 | 0.157 | 1.130 | 0.954-1.339 |
| SR production type | Meat | Ref |  |  |  |  |  |
|  | Multipurpose | 0.140 | 0.066 | 4.505 | 0.034 | 1.150 | 1.011-1.309 |
|  | Mixed | -0.355 | 0.147 | 5.802 | 0.016 | 0.701 | 0.525-0.936 |
|  | Dairy | 0.707 | 0.236 | 8.968 | 0.003 | 2.029 | 1.277-3.224 |
|  | Unidentified | 0.828 | 0.200 | 17.193 | 0.000 | 2.289 | 1.548-3.386 |
| SR housing | Enclosed at night | Ref |  |  |  |  |  |
|  | None | -0.014 | 0.095 | 0.021 | 0.885 | 0.986 | 0.818-1.189 |
|  | Enclosed day and night | -0.664 | 0.205 | 10.490 | 0.001 | 0.515 | 0.344-0.769 |
| SR grazing | Communal | Ref |  |  |  |  |  |
|  | Fenced | -0.117 | 0.099 | 1.408 | 0.235 | 0.889 | 0.733-1.079 |
|  | Mixed | -0.020 | 0.105 | 0.038 | 0.845 | 0.980 | 0.798-1.203 |
|  | Migratory | -0.214 | 0.108 | 3.963 | 0.047 | 0.807 | 0.654-0.997 |
|  | Unidentified | -0.736 | 0.367 | 4.015 | 0.045 | 0.479 | 0.233-0.984 |
|  | Zero-grazing | -0.375 | 0.234 | 2.557 | 0.110 | 0.688 | 0.434-1.088 |
| SR watering | Shared | Ref |  |  |  |  |  |
|  | On-farm | -0.323 | 0.103 | 9.860 | 0.002 | 0.724 | 0.592-0.886 |
|  | Unidentified | 0.120 | 0.238 | 0.255 | 0.613 | 1.128 | 0.707-1.798 |
|  | Mixed | -0.328 | 0.267 | 1.517 | 0.218 | 0.720 | 0.427-1.214 |
| SR breeding method | Own male | Ref |  |  |  |  |  |
|  | Mixed | -.163 | .105 | 2.416 | 0.120 | 0.850 | 0.692-1.043 |
|  | Common-use-male | -.030 | .108 | .079 | 0.779 | 0.970 | 0.786-1.198 |
|  | Unidentified | -.126 | .181 | .484 | 0.487 | 0.882 | 0.619-1.257 |
|  | Male-from-another-farm | -.325 | .150 | 4.716 | 0.030 | 0.723 | 0.539-0.969 |
|  | Artificial insemination | -1.031 | .386 | 7.141 | 0.008 | 0.357 | 0.167-0.760 |

Only multipurpose production type (OR=1.150; 95%CI: 1.011-1.309; p=0.034) and dairy production type (OR=2.029; 95%CI: 1.277-3.224; p=0.003) showed statistically significant positive association when compared with meat production type. Thus multipurpose production type and dairy production type were 1.150 times and 2.029 times more likely to be associated with FMD sero-positivity respectively when compare with meat production type.

Interpretation of OR for risk factors that were negatively associated with FMD sero-positivity was after finding the inverse of OR (1/OR) as specified by Bland and Altman [62]. Therefore, regarding individual animal characteristics, with reference to female animals, male animals were 1.168 times less likely to be seropositive for FMD (OR=0.856; 95%CI: 0.747-0.982; p=0.026). Compared to mature animals, young animals were 1.664 times less likely to be seropositive for FMD (OR=0.601; 95%CI: 0.373-0.970; p=0.037).

With regard to herd level characteristics, animals in the sedentary zone were 2.123 times less likely to be seropositive when compared with those in the pastoral zone (OR=0.471; 95%CI: 0.396-0.559; p<0.001). Unexpectedly, animals in herds that brought in SR were 1.193 times less likely to be seropositive in comparison to animals from herds that did not (OR=0.838; 95%CI: 0.742-0.947; p=0.004). Unexpectedly too and unlike in the bivariable regression model, animals in herds where SR were purchased from the market of middlemen were 1.140 times less likely to be seropositive as opposed to animals from herds where such purchases were not made (OR=0.877; 95%CI: 0.769-0.999; p=0.049). Compared to animals from herds which had wildlife interaction, animals from herds which did not have such interaction were 1.522 times less likely to be seropositive (OR=0.657; 95%CI: 0.535-0.808; p<0.001). Animals under mixed production type when compared with those under meat production type were 1.427 times less likely to be seropositive (OR=0.701; 95%CI: 0.525-0.936; p=0.016). In comparison to animals that were enclosed only at night, animals that were enclosed by day and night were 1.941 times less likely to be seropositive (OR=0.515; 95%CI:0.344-0.769; p=0.001. Migratory animals were 1.239 times less likely to be seropositive when compared with communally grazed animals (OR=0.807; 95%CI: 0.654-0.997; p=0.047. Animals under on-farm watering system were 1.383 times less likely to be seropositive as compared to animals under shared watering system (OR=0.724; 95%CI: 0.592-0.886; p=0.002). Compared to animals under own-male breeding method, animals under male-from-another-farm breeding method were 1.383 times less likely to be seropositive (OR=0.723; 95%CI: 0.539-0.969; p=0.030 while those that were under artificial insemination were 2.801 times less likely to be seropositive (OR=0.357; 95%CI: 0.167-0.760; p=0.008).

## Discussion

The mean SR herd sizes in the PZ and SZ were 27.5 and 2.7 respectively. This is consistent with what has been reported in sub-saharan sedentary production systems [68]. The herd structure in the pastoral zone is similar to what has been reported in Somalia [69]. For the PZ, this is within the range reported recently in Kenya [25] but lower than that reported by Zaal [23], probably due to dwindling land available for livestock keeping and other changes in farming systems. It is true that according to this study, the bulk of SR in Kenya are held in the PZ.

In this study the country sero-prevalence of FMD in SR was found to be 23.3% similar to what has been reported in other countries where FMD is endemic [41,42]. It is however higher than that reported in Ethiopia, Israel, Libya and Sudan [33,34,36,37,43] but about half of what has been reported in Tanzania and Myanmar [39,44,45]. A previous study in cattle in Kenya showed much higher sero-prevalence in cattle at 52.5% [70] and unpublished data obtained at the same time with this current study in Kenyan cattle revealed a sero-prevalence rate of 37.6% in cattle. This means sheep and goats in Kenya are less susceptible to FMDV compared to cattle despite the fact that they are normally herded together in endemic settings of Kenya. The same was observed in Ethiopia [33,34].

The sero-prevalence was significantly higher in the PZ (31.5%) than in the SZ (14.5%).This may be attributed to a high level of herd mobility, contact of animals at grazing and watering points, dynamism of herds (frequent additions) and frequent contact with the livestock of neighbouring countries through cross-border contact in the PZ. These animals move across the boundaries for grazing, watering and also for illegal trade thus promoting the concept that FMD outbreak peaks is associated with animal movement. In the process of movement they also come in contact with other animals from different areas which are an important factor for the transmission of the disease. The livestock in pastoral areas also end up in some sedentary zones during the dry season, potentially spreading disease [71]. Foot and mouth being a disease spread mainly by contact makes sedentary zone have lower incidence of spread between herds. This is important because most of the SR are in the PZ where they are more often herded together with cattle and therefore pose a significant risk. The sero-prevalence in counties within the PZ or bordering the PZ such as Lamu were significantly higher than those in the counties in the SZ as also reported by others [70,72]. This might be due to differences in the movement and distribution of livestock, the level of contact between herds and ungulate wildlife, proximity to stock routes, the grazing patterns and watering sources in each county.

At individual animal level, caprine (goats) showed higher seroprevalence than ovine (sheep). This is in agreement with the findings by Mesfine et al. [33] but in contrast to what has been reported by other researchers who found FMD sero-positivity to be significantly higher in sheep than in goats [38,40,46]. Others found sero-prevalence to be significantly higher in goats than in sheep [35,42]. Exotic breeds had significantly lower sero-positivity (17.7%) compared to both local (24.4%) and cross-breed (24.0%) sheep and goats. This was likely to be as a result of less exposure than susceptibility. Most exotic dairy/meat goats and sheep are usually kept in the intensive production system where enclosures are common therefore contact with other animals is minimal or absent. On the other hand cross breeds and local breeds are more common in the marginal areas where movement and herd interaction are high. A significant difference was observed in sero-prevalence of FMD among mature (23.6%) and young sheep and goats (10.1%). This is in agreement with the results of others [44,45] although the sero-positivity levels in our study were lower. The difference in sero-positivity between age groups may be due to the fact that mature animals may have experienced more exposures to FMD at grazing, watering point and at market than in age group less than one year. Therefore, adult animals might have acquired infection from multiple strains and serotypes thus producing antibodies against multiple virus incursions of FMD. The low prevalence in young animals may also be indicative of persistent passive immunity and less frequency of exposure of the animal to the disease as the farmers keep their lambs and kids in the homesteads. Conversely, there was no significant difference in sero-positivity between sexes though females posted higher seroprevalence rates at 23.8% than males (21.9 %). Similar studies in Kenya and neighbouring Ethiopia reported similar results of no difference in sero-positivity between the sexes [72,73]. However, these results are in contrast to Ethiopian studies where 15.7% and 8.3% seroconversions were reported in male and female animals respectively [74] and 8.9% in female while 3.0% in male [33].The sero-prevalence in animals that were born in the herd was significantly higher than that of animals that were brought in. This is an indication that the FMDV may be circulating in a significant proportion of closed SR herds.

At herd level, unexpectedly, animals from herds that brought in SR had lower sero-prevalence than those that did not. The likely position is that most of those brought in were for breeding purposes from clean farms where good biosecurity is often observed and few could have been from markets and exposed farms. However, in Tanzania it has been shown that herds that recently acquired new livestock had higher sero-prevalence [45]. Further, in our study, animals from herds in which SR were purchased from the markets and middlemen had higher sero-prevalence than those from herds in which purchase was contrary. This stresses the need for purchase of SR from clean sources to avoid introduction of FMD infection in herds. Interaction of herds with wildlife resulted in significantly higher sero-prevalence than when there was no such interaction. Casey-Bryars [45] reported that contact with other FMD susceptible wildlife did not increase the likelihood of FMD in livestock. However in a large proportion of sero-positive herds, it was not clear whether such interaction existed (onknown). This calls for further investigation into the role of wildlife in FMD sero-prevalence in SR.

Sero-prevalence was significantly higher in the multi-purpose production type than in all the other production types (meat, mixed, dairy). This may be possible because this production type is found among pastoral and agro-pastoral systems where purchase of animals is from the market or middlemen and which in each case had high sero-prevalence. Elnekave et al. [36] have demonstrated higher FMD sero-prevalence in dairy animals than in feedlot animals. That sero-prevalence in herds in pastoral and agro-pastoral systems is higher than in other production systems has also been demonstrated in Tanzania [45]. However, Mesfine et al. [33] in Ethiopia have demonstrated the contrary. Further, there was high sero-prevalence in animals whose production type was not identified hence the need for further investigation of sero-prevalence between the production types.

Herds in which animals were not enclosed or enclosed only at night had a significantly higher sero-positivity than herds which were enclosed by day and by night. This could be because enclosure limits mobility and mixing with other herds and wildlife thus reducing exposure to FMDV. Sero-prevalence among animals in herds which had communal, mixed or migratory grazing systems was higher compared to those within fenced or zero-grazing systems. Elnekave et al. [36] have demonstrated that grazing animals compared to feedlot ones are at higher risk of FMD infection. This is expected as in the former three, there is high mobility of animals and therefore high probability of mixing with infected herds. Small ruminants with shared watering showed higher sero-positivity compared with those with on-farm and mixed methods of watering. Sharing of watering creates greater opportunity for animals to mix and therefore be exposed to FMDV than in the other two watering systems.

Animals in herds where own-male, mixed or common-use-male breeding methods were practiced registered significantly higher sero-prevalence than those in herds where male-from-another farm and artificial insemination were the breeding methods used. The high sero-prevalence in own-male breeding method herds further indicates that FMDV may be circulating in closed herds. Mixed and common-use-male breeding methods may expose animals to FMDV through contact with other herds as this type of breeding males which are more in the PZ are usually left to run with the rest of the herd. Usually there are contact restrictions when a male is borrowed from a specific farm and contact is minimal in artificial insemination. Sero-prevalence was higher in low lying areas than in highland areas. This could be because most of the animals in the lowlands are under pastoral and agro-pastoral production systems where sero-prevalence is also high.

After controlling for confounding using multivariable regression analysis, only production type showed a statistically significant positive association with FMD sero-positivity. Sex, age, production zone, bringing in of SR, purchase of SR from the market and middlemen, no wildlife interaction, no housing, grazing, watering and breeding method showed a statistically significant negative relationship. The observation that animals from dairy production type were positively associated with FMD sero-positivity was unexpected but consistent with the findings of Elnekave et al. [36]. Similarly the findings that bringing in of animals, purchase of animals from markets and middlemen as well as migratory grazing system were negatively associated with FMD sero-positivity were unexpected. These unexpected findings may further support the argument that FMD may be circulating in a significant proportion of closed SR herds. The other results are consistent with the findings of this study on FMD sero-positivity per individual and herd risk factors.

It is worth noting that most husbandry related variables showed significant relationship with sero-positivity as has also been alluded to by Balinda et al. [46] in Uganda. These results demand for risk-based surveillance which considers the significant risk factors. They also call for extension services and policies for small ruminant keepers to advice on interventions and husbandry practices which could limit the circulation of FMDV among SR herds which could also reduce cross-contamination with cattle herds. The SR population totaling 44.8 million in 2009 (27.7 million goats and 17.1 million sheep) in Kenya [28] is much higher than large ruminant population (17.5million), therefore infection of SR in sub-clinical levels can have a major impact in transmission to cattle which are usually herded together.

Vaccination of small ruminants against FMD in Kenya is non-existent due to scarce vaccine and cost implications [49]. It may be worthwhile to vaccinate SR in some scenarios, given the identified risk factors. The possibility of transmission of FMDV from cattle to SR needs to be researched. These current Kenyan results have added onto the growing body of literature in Kenya and in the region to demonstrate the significance of SR in the epidemiology of FMD in endemic settings towards better FMD control. But then, there was significant sero-positivity in animals where some variable levels were unidentified hence the need to investigate further the level of sero-positivity with regard to these variables. The results in some counties are useful in making conclusions about the status of FMD in SR and formulating control strategies. However, in some counties the sample size was too small to make any meaningful conclusions and therefore planned studies with sufficient sample sizes are required. More risk factors should be identified through planned studies.

## Conclusion

The bulk of small ruminants in Kenya are held in the pastoral zone. The findings of this study give an understanding of the potential role of small ruminants in the epidemiology of FMD in Kenya and contribute to the global scenario. The study has further established the endemicity of FMD in small domestic ruminants in Kenya. It has shown that the sero-prevalence in small ruminants in Kenya is estimated at 23.3% with sheep and goats having almost equal sero-prevalence. Though sero-prevalence is lower in SR than that already reported in cattle, the revealed dynamics mean SR are potential transmitters and maintenance hosts of the disease in cattle. Cross contamination with FMDV across the species needs investigation. The pastoral zone had higher sero-positivity as compared to sedentary zones. This outlays the importance of concentrating control efforts in the pastoral zone where sero-positivity is high without neglecting the sedentary areas which post the highest productivity losses in case of FMD outbreaks. Besides, livestock in the pastoral zone also end up in some counties in the sedentary zone during the dry season, potentially spreading disease.

Risk factors for sero-positivity in this study were mainly husbandry related. There is sufficient evidence that the FMDV may be circulating in a significant proportion of closed SR herds given the near ubiquitous distribution of the disease and the negative associations of sero-positivity with risk factors related to introduction of animals into the herds. Past efforts for control of FMD in Kenya centered on what was commonly referred to as compulsory vaccination areas mostly located in the sedentary areas. The findings of this study should be considered in the development of FMD risk-based surveillance and control plans in small ruminants alongside those of the large ruminant population.

## Acknowledgement

This study had support from a project entitled “Improving Animal Disease Surveillance in Support of Trade in IGAD Member States”, in short “Surveillance of Trade Sensitive Diseases – STSD”. This was a regional component of the Supporting the Horn of Africa’s Resilience (SHARE). The project was implemented by IGAD member states through AU-IBAR and IGAD. The ELISA Kits used in this work were provided by Eu-FMD through the Nakuru FMD Real - Time Training Course credit points We acknowledge approval of the study by the Directorate of Veterinary Services (DVS), Kenya and respective County governments. The survey teams and Foot- and -mouth Disease Laboratory staff are acknowledged for sample and data collection as well as sample testing. The respective County livestock keepers are acknowledged for presentation of animals and provision of data. The DVS data entry team comprised of Ruth Manasse, Peninah Khan and Nelly Achieng’ are also acknowledged.

## Supporting information

**S1 Table. Descriptive statistics of sampled individual animal variables in the pastoral and sedentary zones, Kenya, 2016**

**S2 Table. Descriptive statistics of herd level variables in the pastoral and sedentary zones, Kenya, 2016**

**S3 Table. Small ruminant FMD Sero-positivity per county, Kenya, 2016**

**S4 Table. Bivariable regression model of individual animal risk factors for FMD Sero-positivity, Kenya, 2016**

**S5 Table. Bivariable regression model of herd level risk factors for FMD sero-positivity in small ruminants, Kenya, 2016**

**S1 File. Data collection questionnaire**

**S2 File. Questionnaire guidelines**

**S3 File. Sample collection form**

**S4 File. Survey data**

**S5. Survey data code book**

